# The influence of movement-related costs when searching to act and acting to search

**DOI:** 10.1101/2022.03.01.482521

**Authors:** J. B. Moskowitz, S. A. Berger, M. S. Castelhano, J. P. Gallivan, J. R. Flanagan

## Abstract

Real world search behaviour often involves limb movements, either during search or following search. Here we investigated whether movement-related costs influence search behaviour in two kinds of search tasks. In our *visual* search tasks, participants made saccades to find a target object among distractors and then moved a cursor, controlled by the handle of a robotic manipulandum, to the target. In our *manual* search tasks, participants moved the cursor to perform the search, placing it onto objects to reveal their identity as either a target or a distractor. Across experiments, we manipulated either the effort or time costs associated with movement such that these costs varied across the search space. We varied effort by applying different resistive forces to the handle and we varied time costs by altering the speed of the cursor. Our analysis of cursor and eye movements during manual and visual search, respectively, showed that effort influenced manual search but did not influence visual search. In contrast, time costs influenced both visual and manual search. Our results demonstrate that, in addition to perceptual and cognitive factors, movement-related costs can also influence search behaviour.

**Public Significance Statement:** Many of the tasks we perform on a daily basis involve searching for targets. Numerous studies have investigated perceptual and cognitive factors that influence decisions about where to search. However, few studies have examined how search is influenced by movement-related costs associated with manual search (e.g., opening drawers to find a corkscrew) or acting on an object once it has been located (e.g., reaching for a particular bottle of wine once it has been spied in a rack). We show that movement effort and time costs associated with manual search, and time costs associated with moving after visual search, can influence decision-making about where to search over time.

## Introduction

Visual search behaviour in humans has been studied extensively, with evidence suggesting that search is driven by both bottom-up (i.e., stimulus-driven) and top-down (i.e., goal-oriented) influences on attention (Castelhano & Henderson, 2007; Corbetta & Shulman, 2002; Desimone & Duncan, 1995; Draschkow & Võ, 2017; Henderson & Hollingworth, 1999; Wolfe & Horowitz, 2017). Most studies of search behaviour have used visual search tasks that involve locating a target item and producing a response (e.g., a button press) once it is located. However, real world search behaviour often involves significant movement, whether moving within and acting on the environment in order to perform the search, or acting on a target object once it has been successfully located (for a review see Hayhoe, 2017; Land et al., 1999; Land & Hayhoe, 2001).

Most research on visual search has used tasks that do not involve acting on (e.g., reaching towards) located targets, and thus has not considered the possible influence of movement related factors on search behaviour. According to serial models of behaviour (McClelland, 1979; Miller et al., 1960; Sternberg, 1969), movement planning only occurs after the decision about where to move is made. Thus, serial models predict that decisions about where to look while visually searching for a reach target are independent of planning the subsequent reach movement, and decisions about where to reach during manual search are likewise independent of reach planning. However, this serial view has been challenged by converging neurophysiological and behavioural evidence indicating that different movement options can be specified or planned prior to deciding which option to execute (Cisek & Kalaska, 2005; Gallivan et al., 2015; Klaes et al., 2011; Pastor-Bernier & Cisek, 2011; Song & Nakayama, 2009; Thura & Cisek, 2014). Planning potential movement options in advance of movement selection may provide a mechanism through which movement-related costs can be factored into decision-making about where and when to move (Cisek, 2007; Cisek & Kalaska, 2010).

Previous work has shown that properties of movements, including variability, duration, and effort, can influence decision making behaviour in sensorimotor tasks (Gallivan et al., 2018; Hayhoe, 2017; Hayhoe & Ballard, 2005). For instance, studies of target-directed reaching have found that people can factor into account their motor variability when making strategic decisions about where and when to reach (Battaglia & Schrater, 2007; Diamond et al., 2017; Faisal & Wolpert, 2009; Moskowitz et al., 2020; Trommershäuser et al., 2003, 2005, 2008). Movement costs, such as energy expenditure, are an integral component of most models of motor control and can influence decisions about how to move in order to achieve a movement goal, including how to respond to feedback during the movement (Harris & Wolpert, 1998; Scott, 2004; Todorov, 2004; Todorov & Jordan, 2002). Movement effort can also influence action selection. In a task where participants could freely choose between two possible reach targets, participants preferred movements to the target associated with less biomechanical effort (Cos et al., 2011, 2012, 2014). Similarly, when walking, people select footholds that minimize energetic costs through the maintenance of a stable gait (Domínguez-Zamora & Marigold, 2019; Matthis et al., 2018).

Recently, studies have shown that movement costs associated with responding are also capable of biasing decision making in perceptual judgement tasks (Hagura et al., 2017; Marcos et al., 2015). Hagura et al. (2017) asked participants to report whether they saw dots in a display moving coherently either to the left or right by moving either the left or right hand, respectively. Participants held a handle in each hand, which applied a resistive load during hand movement. When the resistive load incurred when moving one of the hands was increased relative to the other hand, perceptual judgments became biased toward the direction associated with the hand that was easier to move. In a similar experiment, participants reported the direction of dot motion by reaching to a target located on the left and right (Burk et al., 2014). After initiating the reach, participants sometimes changed their mind, based on visual evidence obtained after the initial decision to move, and reversed their reach direction. It was found that participants were less likely to change their mind when the two targets were far apart, such that greater effort was required to correct the movement. Together, this work suggests that, when relaying a decision via arm movements, costs incurred at the output-level of the motor system can seemingly influence processes occurring at the level of the visual-perceptual system.

A handful of studies have shown that people are more likely to use memory to guide search when search involves more effortful movement (Ballard et al., 1995; Gilchrist et al., 2001; Kit et al., 2014; Li et al., 2016, 2018; Smith et al., 2008; Solman & Kingstone, 2014). Solman and Kingstone (2014) examined a search task in which participants viewed items with different letters on them, and where the target letter was varied from trial to trial. The locations of the items were either randomized on each trial or repeated across trials. In addition, the size of the display was varied such that search either required both eye and head movement or only eye movements. The authors found that the reduction in search time between randomized and repeated displays was greater when search required both eye and head movements. This suggests that when search required head movements, participants exploited memory of the items to a greater extent (Solman & Kingstone, 2014). Recent studies examining search in virtual environments, in which participants walk around, have also reported that participants rely more strongly on memory than in standard laboratory search tasks (Kit et al., 2014; Li et al., 2016, 2018). Ballard and colleagues (1995) also showed that costs associated with gaze shifts can influence the contribution of memory in a task in which participants had to arrange a set of blocks to match a visible model showing the desired arrangement. They found that participants fixated the model less frequently when gaze shifts, between the model and the set of blocks, required more costly head movement in comparison to when they only required less costly eye movement. Together, these studies demonstrate how increased effort leads to an increased influence of memory on search performance and decision making.

### Present Study

To our knowledge, previous research has not directly examined whether visual or manual search can be influenced by movement costs associated with reaching towards or locating a target object. To investigate this issue, we designed a series of experiments where participants searched a display containing target and distractor objects and incurred movement-related costs when either moving a cursor to a target object once it is located (visual search), or by moving a cursor to objects in the scene in order to reveal whether they are a target or a distractor (manual search). We separately examined effort and time costs associated with movement. To assess effort costs we applied forces to the hand through the handle of a robot manipulandum that the participant moved in order to control the position of the cursor. To assess time costs we had participants move the cursor with a joystick and manipulated the speed of the cursor. These costs were always on a spatial gradient, such that the cost of moving depended on the spatial location of the cursor in the search space. In our search tasks, multiple potential target objects were presented in each trial and randomly distributed in the display, meaning that participants would still have a high probability of successfully locating a target object regardless of whether they searched in a high or low movement cost location. If the costs of movement are factored into search decisions, we would expect to see a shift in search behaviour, with search being directed towards locations that reduce movement effort or duration.

In Experiment 1 we tested whether movement effort influences visual search using a ‘search-and-then-reach’ task in which participants were asked to visually search for a target object among distractors, and then reach for the target using a cursor controlled by the handle of a robotic manipulandum. Participants were required to visually locate one of two targets, and then move a cursor from the center of the display onto the target object. The target and distractor objects were designed such that identifying the target object required foveal vision. Therefore, we could use eye movements to determine where a participant was searching for the target. We manipulated the effort associated with reaching to the target by applying a large resistive, viscous (i.e. velocity-dependent) force to the handle when it moved on either the left or right side of the search space (counterbalanced across participants). We predicted that participants would avoid searching the side of space associated with greater movement costs (i.e., greater viscosity).

The aim of Experiment 2 was to examine whether, and if so how, effort costs that are incurred *during* search influence search behaviour. In this experiment, participants performed an ‘act-to-search’ task in which hand movements were required to perform the search. Participants moved the handle of a robotic manipulandum to move the cursor to objects in a display in order to reveal the identity of the object (target or distractor). If the revealed object was a target, the trial ended, otherwise participants had to continue their search. We applied an elastic force to the handle that was proportional to its distance from the start position such that greater effort was required to place the cursor on objects located farther from the start. Across blocks of trials, participants searched for a target object with the elastic force turned either on or off. We predicted that when the elastic force was on, participants would visit, on average, objects closer to the start location in comparison to when the elastic force was off.

In Experiments 3 and 4 we tested whether movement time costs influence manual and visual search, respectively. The manual search task involved moving a cursor to an object to reveal its identity (target or distractor) and the visual search task required foveating an object to determine its identity. In the manual search task performed in Experiment 3, cursor movement was controlled by a joystick. We manipulated the time required to move in different regions of the search space by modifying the speed of the cursor based on its radial angle from the start position, with the cursor moving faster when it was located on either the left or right side of the search space (counterbalanced across blocks). We predicted that participants would more often visit objects (to determine whether the object is a target) on the side of space associated with faster cursor speeds. In Experiment 4, participants performed a block of trials in the manual search task with the cursor moving faster on one side of the search space, and then completed a block of visual search trials with the same cursor speed mapping. Participants then performed two additional blocks of the manual and visual search tasks with the cursor moving faster on the other side of the search space. We included these manual search trials to ensure that participants understood how time costs varied across the search space. We predicted that in visual search trials, search (as measured by gaze) would be biased to the side with the faster cursor movements.

### Experiment 1

In Experiment 1 we tested the novel hypothesis that the physical effort associated with reaching a target object, once it has been located among distractors, can influence the preceding visual search behaviour as measured by gaze. Participants made reaching movements with a cursor controlled by a grasped manipulandum (Figure 1A). We applied a large resistive force to the handle of the manipulandum when participants moved on one side of the search space (counterbalanced across participants) but not the other. We predicted that participants would quickly learn the association between effort and spatial location across trials, and then bias their visual search to the low effort side. The display viewed by participants always contained 30 objects, with 15 on either side of midline (Figure 1B). Two of these objects were targets and these were randomly located such that at least one target was located on a given side on 75 percent of the trials. Thus, in principle, participants could reduce the overall effort required to perform the task by first searching the lower-effort side of space.

**Figure 1.**
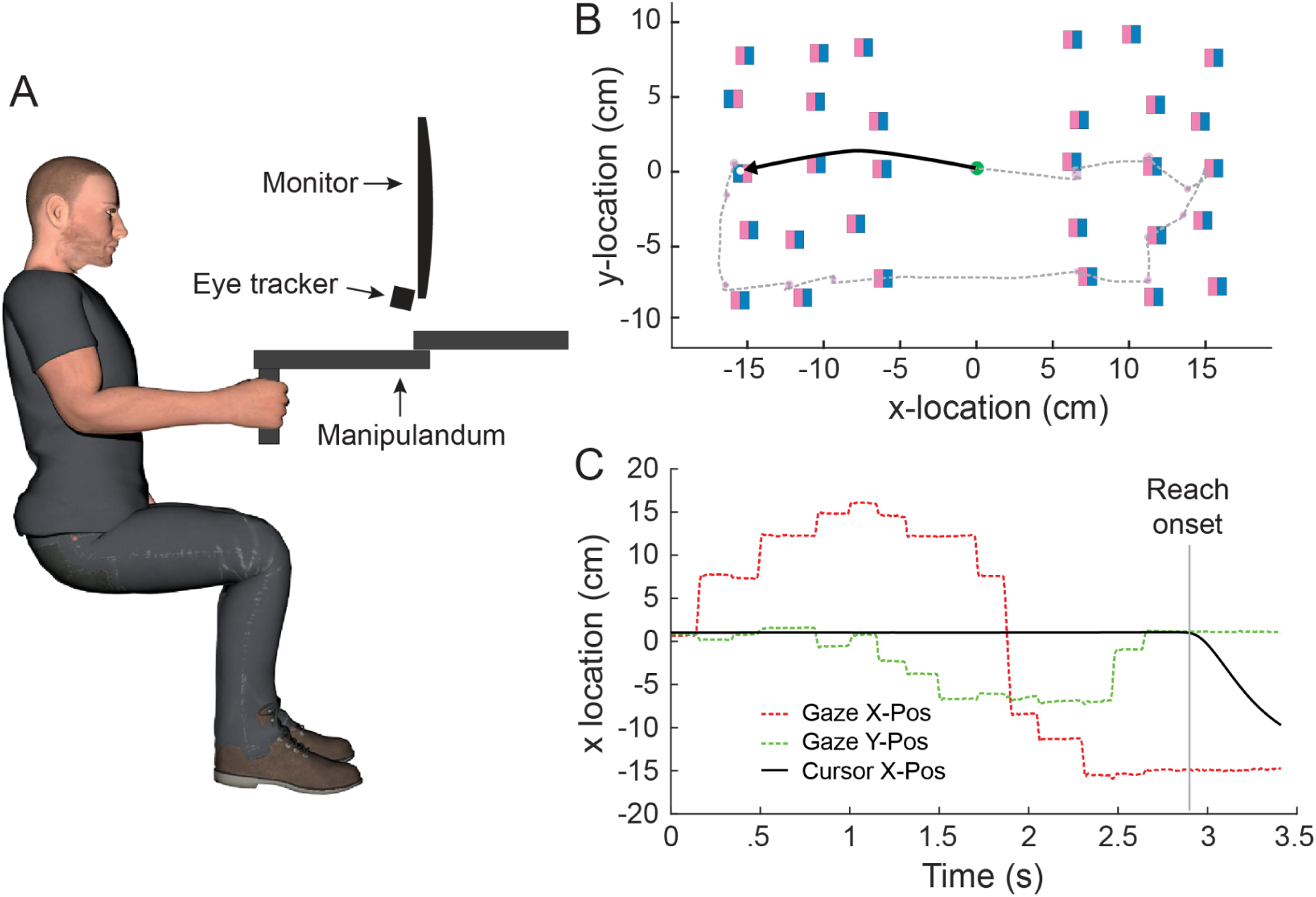
Experimental set up and data from an exemplar trial. A) Participants moved a cursor to target objects located on a vertical screen by moving the grasped handle of a robotic manipulandum in the horizontal plane. The manipulandum could apply viscous forces to the hand during movement. Gaze was recorded with an infrared video-based eye tracker. B) In each trial 30 objects were presented on the screen, with 15 on either side of midline. The 30 objects included two target objects (pink on the right side) and 28 distractor objects (pink on the left side). The dashed and solid traces show the gaze and cursor paths, respectively, for a single baseline trial in which equal and small viscous forces were applied to movements on the left and right sides of the search space. Gaze and the cursor were at the start position (green circle) at the start of the trial. C) Time varying X and Y gaze locations and the X position of the cursor for the trial shown in B.

## Methods

### Participants

Eleven participants (8 female) between the ages of 18 and 22 years old (M = 19.6) completed the experiment. Three additional participants who were recruited did not complete the experiment because we were unable to achieve an adequate gaze calibration. Participants were required to be right handed and have normal vision or corrected-to-normal vision while wearing contacts. All participants were compensated $15 or 1.0 course credits for their participation. Participants provided written informed consent and after the conclusion of the experiment they were debriefed. The experiment was approved by the Queen’s General Research Ethics Board and complied with the Declaration of Helsinki.

Although we are unaware of any studies that have examined how costs associated with manual action influence visual search behaviour, many previous studies of motor learning have examined how such costs affect decision-making and behaviour (e.g., Gallivan et al., 2018; Shadmehr, Smith, et al., 2010; Wolpert & Landy, 2012). In most motor learning experiments, there are between eight and twelve participants per experimental group. This sample size provides sufficient power to detect the large effects typical of motor learning experiments, where the effect of interest is observed in most, if not all, participants. In a previous study (Diamond et al., 2017), we examined how motor costs influence decision-making in a target foraging task, in which different groups of participants harvested targets with either hand or eye movements. In that study, we found an effect size of Cohen’s d = 1.1. A power analysis using G*Power 3.1 (Faul et al., 2009) indicated that with this effect size and a power of 0.90, 11 participants would be required. Therefore, we aimed to obtain at least 11 participants in each experimental group.

#### Apparatus and stimuli

Seated participants used their dominant hand to grasp the handle of a planar robotic manipulandum (Figure 1A; Kinarm End-Point, Kinarm, Kingston, ON, Canada) and viewed the visual stimuli (Figure 1B)—including the target objects, distractor objects, and a cursor controlled by handle movement—on a vertical monitor positioned directly in front of them. The position of the cursor (filled white circle, radius 3 mm) on the monitor was linked to the position of the handle in a horizontal plane. The direction mapping between handle and cursor movement was the same as a standard computer mouse, such that forward and backward movements of the handle moved the cursor up and down, and right and left handle movements moved the cursor right and left. When the cursor was in the center of the screen, the handle was located ∼20 cm in front of the participant’s chest and in the mid-sagittal plane. There was a 1:1 correspondence between the distance moved by the handle in the horizontal plane and the distance moved by the cursor on the screen. The position and velocity of the handle and the programmed resistive force it applied to the hand were recorded at 1000 Hz. Gaze data were collected at a rate of 500 Hz using an infrared eye tracker (Eyelink 1000, SR Research, Ottawa, ON, Canada) mounted just below the display monitor. A chin rest (not shown in Figure 1A) was used to limit head motion during the experiment.

At the beginning of each trial, a start position (empty green circle, radius 5 mm) appeared at the center of the monitor. Once participants moved the cursor to this location, it filled solid green, and after a delay of 750 ms, a fixation cross appeared over it (solid white, width 1.4 cm). Participants were instructed to fixate the cross for 1000 ms at which point the target and distractor objects appeared. In all trials, there were 28 distractor objects and 2 target objects, with 15 objects on each side located in cells of a 5 × 3 grid (Figure 1B). The size of each cell of the grid was 4 × 4 cm and the position of each object within the cell was randomly jittered. The objects were 1.2 cm wide squares (subtending ∼1.9º of visual angle when in the center of the monitor). For the target objects, the right half was coloured pink and the left half coloured blue. The distractor objects had the opposite colour arrangement. The locations of the target objects (i.e., the cells in which the targets appeared) were pseudo-randomized such that each target appeared an equal number of times in each cell and a given cell could not contain a target over successive trials. Over the course of the experiment, both targets were on the left side in 25% of the trials, both targets were on the right side in 25% of the trials, and there was one target on either side in 50% of the trials. The participant was required to find one of the two target objects and then move the cursor to that target. The trial was considered to have been completed when any part of the cursor overlapped with any part of the target for 100 ms.

A movement time criterion was imposed such that if participants took longer than 2 s to reach the target once they initiated the movement, they would be presented with the phrase “TOO SLOW” in the center of the display and hear an ‘incorrect’ tone (5 Hz, 100 ms) being played. We included this movement time criterion to ensure that participants always experienced fairly significant resistance when velocity-dependent forces were imposed by the handle (see below). To avoid excessive search times, we also implemented a combined search plus movement time criterion such that if the target was not reached within 10 seconds, participants were presented with the phrase “TIMEOUT” and the same incorrect tone. This time limit was exceeded in only 1% of all trials. If participants completed the trial within these time criteria, they were presented with the phrase “TARGET FOUND” and a ‘correct’ tone (5000 Hz, 100 ms) was played.

#### Procedure

Prior to beginning the experiment, participants completed an eye calibration procedure followed by five practice trials. The experiment started with 30 baseline trials in which a small viscous (i.e. velocity-dependent) load of 10 Ns/m was applied to the handle when reaching to targets on either the left or right side. Note that all viscous forces were resistive and acted in the opposite direction of the motion of the cursor. By including these baseline trials, we could measure each participant’s initial bias in search behaviour.

Following the baseline trials, participants completed 180 test trials, taking a short rest every 60 trials. In the test trials, we set the viscosity of the load on the higher effort side to 30 Ns/m while keeping the low effort side at the baseline value of 10 Ns/m. Figure 2 shows peak force applied to the hand and peak hand velocity for movements to targets on the low and high effort sides of space. As expected, peak forces were far greater for movements on the high effort side (*t*(10) = 22.56, *p* < .001). Peak velocity was slightly but significantly smaller for movements on the high effort side (*t*(10) = 5.82, *p* < .001). After the test phase, participants performed an additional block of 30 baseline trials, with viscosity returning to 10 Ns/m on both sides, which we refer to as the washout phase. After all trials had been completed, we asked participants a series of questions to gauge their understanding of the experiment. Specifically, we asked, in order, if they noticed any changes during the experiment; if they noticed a resistance during movement; if they noticed whether the resistance increased at any point during the experiment; and finally, whether that increase was tied to movement to either the left or right side of the workspace.

**Figure 2.**
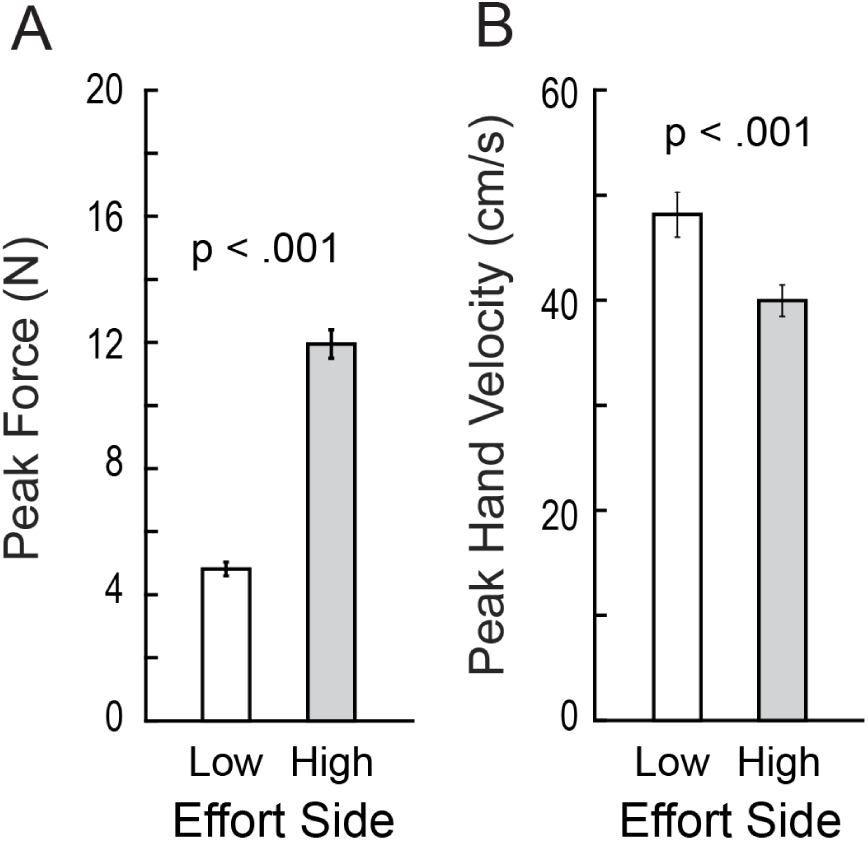
Participants’ movements required more force generation, and were slower, on the high effort side of space. (A) Peak forces applied to the hand by the handle during reaches to target objects located on the high and low effort sides of the search space. (B) Corresponding peak hand velocities of these reach movements. Error bars indicate ±1 standard error.

#### Data Analysis

After eliminating blinks, the raw gaze signal was smoothed using a second-order, zero-phase lag Butterworth filter with a cutoff frequency of 50 Hz. We then extracted the fixation locations for each trial from the time the objects were presented until reach onset (i.e., the time at which hand speed exceeded 5 cm/s), excluding the first fixation location centered on the fixation cross. For each trial, we attempted to assign each fixation to one of the 30 objects. Specifically, we assigned each fixation to the closest object, provided it was no more than 2 cm in distance from the center of that object. Less than 1% of all fixations could not be assigned to an object.

## Results and Discussion

Figure 1B shows the gaze and cursor paths for a single baseline trial from one of our participants. Figure 1C shows the x and y gaze positions, and the x cursor position, as a function of time for the same trial. In this trial, the search time (from the onset of the search stimuli to the onset of the reaching movement) was 3.8 seconds, during which 10 objects, including one of the targets, were fixated. Across participants, the average trial search time, based on participant means, was 2.8 s (SE = .09 s) and the corresponding average number of objects fixated was 5.8 (SE = .3).

Given our hypothesis that increased motor effort would bias search behaviour, we were primarily interested in which side of the search space participants directed their gaze during search. Therefore, for each trial, we first computed the average x location of each fixation prior to reach onset, where the x location of the center of the search space is zero (Figure 1B). We then signed this location as positive or negative depending on whether it was on the low effort or high effort side, respectively, and multiplied it by the duration of the fixation. We then summed up these values, across the fixations in the trial, and divided by the total fixation duration in the trial in order to normalize across trials of varying search duration. We refer to this measure as the ‘integrated gaze location’, with positive values indicating that gaze was biased towards that low effort side (i.e., the side requiring less effort in the test phase) and negative values indicating that gaze was biased towards that high effort side. Note that when computing the integrated gaze location in baseline trials, we used the low and high effort sides from the (later) test phase experienced by the participant, allowing us to remove any baseline bias. Other assessed other measures to evaluate gaze bias, including the proportion of fixations on the force minimum side and the proportion of time spent fixating the force minimum side. Because all of these measures revealed very similar patterns of results, we opted to only report the results for the integrated gaze location.

Figure 3 shows the relationship, across participants, between the average integrated fixation location in baseline and test trials. Each circle represents a single participant and, as noted above, positive values indicate a bias to searching on the low effort side. Participants above the unity line (x=y) searched more on the low effort side during test trials than during baseline trials, as predicted by our hypothesis. Participants along the line did not change their search behaviour from baseline to the test phase, and participants below the line were more biased towards the high effort side during test trials in comparison to baseline trials. Most participants were close to the unity line and, across participants, the average integrated fixation locations in the baseline and test trials were highly correlated, *r* = .92, *p* < .001, indicating that participants generally did not alter their search behaviour from the baseline phase to the test phase.

**Figure 3.**
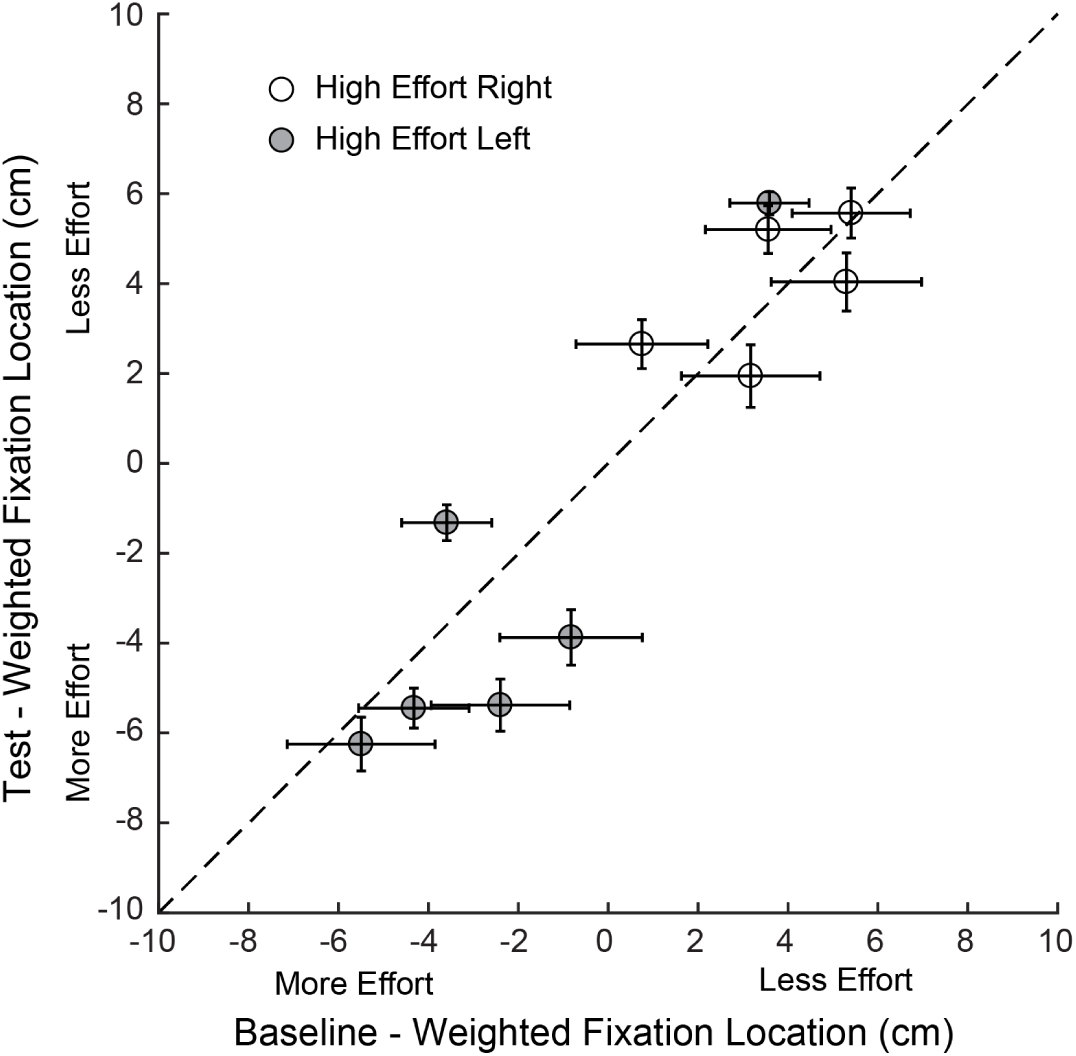
Integrated fixation location in the test phase plotted against the integrated fixation location in the baseline phase. Each circle represents the mean of a single participant and the error bars represent ±1 standard error. Filled and open circles indicate that the location of the high effort side was on the left and right, respectively. Positive values indicate that the integrated fixation location was on the low effort side of the search space. Values located above the dashed unity line indicate that the participant shifted their search towards the low effort side during the test phase.

Figure 4A shows the mean integrated fixation location, averaged across participants, during the baseline, test and washout phases. For each phase, separate bars are shown for each location (left or right) of the low effort side. Overall a clear left side bias was observed such that participants tended to search on the low effort side when the low effort side was on the left (open bars), and on the high effort side when the high effort side was on the left (filled bars). A phase (baseline, test, washout) x high effort side (left, right) mixed model analysis of variance (ANOVA) revealed an effect of high effort side, *F*(1,9) = 10.94, *p* = .009, η^2^ = .549, on integrated fixation location but failed to reveal an effect of phase, *F*(2,18) = .104, *p* = .903, or an interaction between high effort side and phase, *F*(2,18) = .451, *p* = .644. Note that our main hypothesis predicted that there would be an effect of phase. However, this effect was not observed.

**Figure 4.**
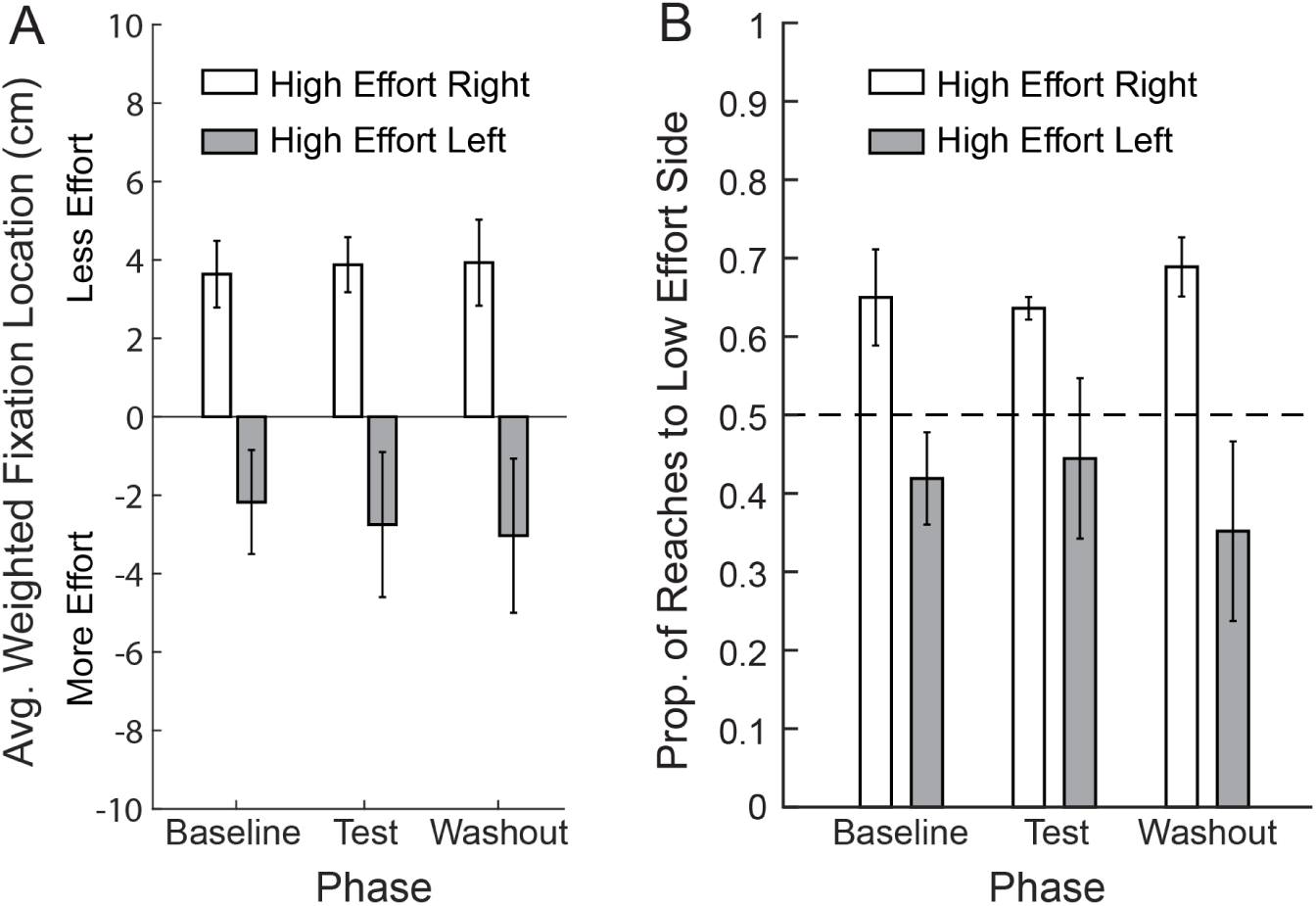
(A) Average integrated fixation location based on participant means. For each phase of the experiment, separate bars are shown for the different load conditions and the location of the high effort side, with positive values indicating fixations were biased to the low effort side. (B) Proportion of reaches to the low effort side during trials where there was one target on each side of the screen. Separate bars are shown for the two load conditions and, for each condition, the location of the high effort side. The dashed horizontal line represents the proportion of reaches expected if participants were selecting a side at random. (A, B) Bars represent participants’ means and the error bars indicate ±1 standard error.

Another way to assess search bias is to examine which target participants reached for in trials in which there was a target on each side. Figure 4B shows the proportion of reaches to the low effort side in each phase (i.e., baseline, test, washout) of the experiment. Separate bars are shown for each location (left or right) of the high effort side. It can be seen that for each high effort side, the proportion of reaches to the low effort side was quite consistent across the phases of the experiment. Consistent with the left side search bias described above, reaches tended to be biased towards the left side of space such that participants tended to reach to the low effort side when the low effort side was on the left (open bars), and on the high effort side when the high effort side was on the left (filled bars). To quantitatively assess this pattern of effects, we performed a phase (baseline, test, washout) x high effort side (left, right) mixed model ANOVA. Consistent with the gaze analysis above, this analysis revealed a main effect of high effort side, *F*(1,9) = 6.713, *p* = .029, η^2^ = .427, but failed to reveal a significant main effect of phase, *F*(2,18) = .097, *p* = .908, or an interaction between high effort side and phase, *F*(2,18) = 1.291, *p* = .299. Again, note that our main hypothesis predicted that there would be an effect of phase.

The results of this initial experiment did not support our hypothesis that visual search behaviour would be influenced by effort costs associated with reaching to targets once they have been located. Specifically, we observed that participants did not tend to alter their search behaviour after forces were introduced that made one side of the search space more effortful to reach in than the other side. A possible explanation for this outcome is that participants selected a side to search early on during the baseline phase and then did not deviate from this strategy. However, many participants did not exhibit a strong bias towards any given side. Another possibility for why movement costs were not factored into account is that they were incurred well after decisions about where to search were made. Recent work on sensorimotor decision making has shown that both effort costs and rewards are temporally discounted, such that their influence diminishes with the delay between when decisions are made and when the costs or rewards are incurred (Berret & Jean, 2016; Rigoux & Guigon, 2012; Shadmehr, Orban de Xivry, et al., 2010). It is also possible our participants simply did not find the load experienced on the handle to be adversive, perhaps because the load was only experienced for a relatively short duration and intermittently.

After the search task had ended, participants were asked a series of questions to gauge their understanding of the forces. Out of the 11 participants, 10 noticed the load on the handle and, of these 10, 6 were able to accurately describe the location of the higher load in the test phase. Based on these qualitative findings, it does not seem that our failure to demonstrate movement cost influences on search can be attributed to a failure of participants to appreciate, at the level of verbal report, that forces were acting on the hand and that these were dependent on reach location.

### Experiment 2

Experiment 1 assessed whether movement costs associated with reaching to a target *after* it was located have an impact on visual search behaviour. The aim of Exp. 2 was to test whether movement costs experienced *during* the act of manually searching influence search behaviour. To this end, we developed a task in which participants manually searched a display of target and distractor objects using a cursor controlled by the handle of the robotic manipulandum. The object was revealed as a target or distractor when the cursor contacted it. In alternating blocks of trials, participants experienced two different load conditions. In force-on trials, a large elastic load was applied to the handle of the manipulandum such that the load linearly increased with the distance of the cursor from the hand start position. In force-off trials, no load was applied. We anticipated that application of this load in force-on trials would cause participants to keep the handle closer to the start position than in force-off trials, thereby reducing the motor effort expended during search. Each participant experienced two force-on blocks and two force-off blocks with the two block types alternating. The initial block type was counterbalanced across participants. We implemented this block structure because of the possibility that any effect of the load on search behaviour might only emerge after experiencing a number of force-on and force-off trials.

We reasoned that movement effort costs might have a greater influence on search behaviour in manual search than in visual search for two reasons. First, in our manual search task, the length of time over which effort costs are experienced is typically greater than in our visual search task (see below). Second, whereas in visual search effort costs are delayed, in manual search they are experienced during search and hence while decisions about where to search are being made.

## Methods

### Participants

Sixteen participants (5 female) between the ages of 18 and 24 years old (M = 19.5) were recruited for this experiment. (See Exp. 1 for sample size considerations.) Participants were required to be right handed, and have normal or corrected-to-normal vision while wearing contacts. All participants were compensated $15 for their participation. Participants provided written informed consent, and after the conclusion of the experiment they were debriefed. The experiment was approved by the Queen’s General Research Ethics Board and complied with the Declaration of Helsinki.

### Apparatus & Stimuli

Participants used the robotic manipulandum described in Exp. 1. Specifically, they controlled a cursor by moving the handle of the manipulandum and there was a 1:1 correspondence between the distance moved by the handle in the horizontal plane and the distance moved by the cursor on the screen. However, in this study, we did not carry out eye tracking, as it was not germain to the main hypothesis under consideration.

The target and distractor objects in this experiment were the same size and appearance as in Exp. 1 and were located within a circular search area that had a radius of 14 cm around the center of the monitor (see Figure 5A). Within this circle, 60 objects were arranged by aligning them to a grid which contained 61 cells, with the extra cell containing the start position (see below). The size of each cell of the grid was 3.5 × 3.5 cm and the position of each object within the cell was randomly jittered.

**Figure 5.**
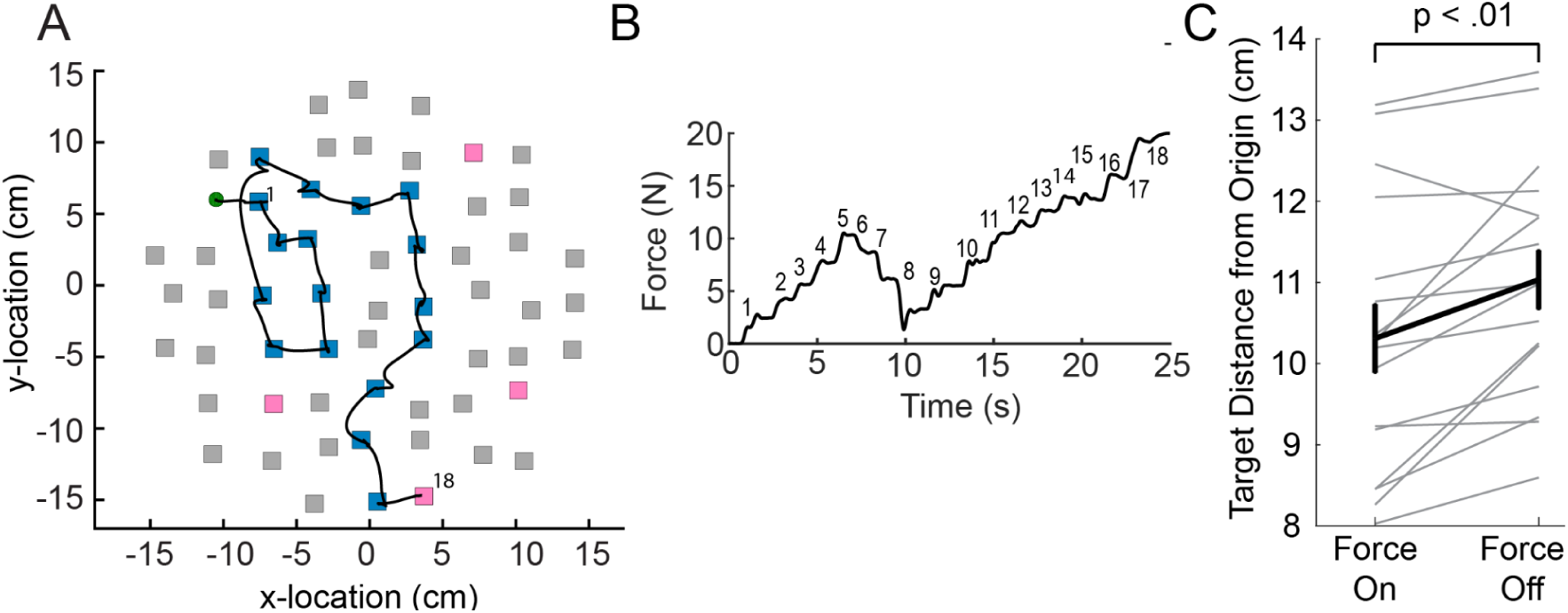
A) In each trial 60 objects were presented on the screen, positioned in a circular grid. The object position within each grid cell was randomly jittered. The 60 objects include four hidden target objects (in pink). The black line shows an example cursor path starting from the origin and ending at the target object at the bottom of the screen. B) Force applied by the manipulandum handle as a function of time in the trial. The times at which the 18 objects visited in this trial are labelled in order. C) Average distance from origin of objects visited across trials for blocks with the elastic force on, and blocks without. Individual participants are shown in gray traces. Error bars indicate ±1 standard error.

For each trial type (i.e., force-on and force-off), the start positions for the first 61 trials were selected by random sampling, without replacement, of all 61 possible target positions. The start positions for the last 39 trials were also selected by random sampling without replacement from a pool of all 61 target positions. For each trial, we randomly sampled without replacement 4 target locations from the 60 possible target locations (i.e., excluding the already selected start position).

As in Exp. 1, participants had to hold the cursor at the start position and were asked to fixate the cross that appeared over it. Once the cursor was held in the start position for 750 ms, the search objects appeared and participants had to locate one of the four search targets. In order to identify whether a given gray search object was a target or a distractor, participants had to bring the cursor to a stop on the object. Specifically, they needed to keep the center of the cursor within 5 mm of the center of the object for 500 ms, after which the search object changed color. If the object was a target, it turned pink, the text “TARGET FOUND” was displayed in the center of the display, a ‘correct’ tone (5000 Hz, 100 ms) sounded, and, after a 1 s delay, the trial ended. However, if the object was a distractor, it turned blue and participants had to continue searching for one of the targets. Once a search object changed colour, it remained that way for the duration of the trial, such that participants did not have to memorize the location of already visited objects. If participants could not locate a target object within 30 s, “TIMEOUT” was displayed on the screen, an incorrect tone (5 Hz, 100 ms) sounded and, after a 1 s delay, the trial ended. This occurred in less than 1% of all trials.

### Procedure

Participants were informed prior to beginning the experiment that there were four search targets on each trial and that their location was determined through randomization. The experimenter demonstrated a trial to familiarize them with the task. Participants completed four blocks of 50 trials each starting with either a force-on or force-off block (counterbalanced across participants) and then alternating between block types, such that all participants experienced two force-on blocks and two force-off blocks. In force-on trials, the manipulandum applied an elastic force to the handle with the force directed back to the start position and increasing linearly with the distance of the cursor from the start position multiplied by the spring constant k = 80 N/m. With the application of this elastic load, the further an object was from the start position, the more effort participants needed to expend to visit it. In force-off trials, no external force was applied to the handle (i.e., k was set to zero).

### Data Analysis

The locations of the search objects visited in each trial were recorded. For each trial, we calculated the distance from each object visited (including the target) to the cursor’s start location. To obtain a measure of how far from the start participants searched, we then computed, for each trial, the average distance of the visited objects from the start position.

## Results and Discussion

Figure 5A shows the cursor path, the objects visited, and the locations of the four targets (shown in pink) for a single trial. Figure 5B showed the force applied by the manipulandum handle as a function of time in this trial. The time at which search objects 1-18 were visited is labelled. On average, the search time across participants was 14.9 s (SE = .4 s) in force-off trials, and 15.3 s (SE = .3 s) in force-on trials. In force-off trials participants visited an average of 11.9 (SE = .3) objects before locating the search target and the average in force-on trials was 11.7 (SE = .2). There was no significant difference between force-on and force-off trials in terms of search time, t(15) = .809, p = .431, or the number of objects visited to locate a search target, t(15) = .433, p = .671.

We predicted that participants would, on average, visit object locations closer to the start position in force-on trials in comparison to force-off trials. To assess this prediction, we determined, for each participant, the mean average distance for each of the four trial blocks. To investigate the effects of block type (force-on versus force-off) and block number (first or second), we ran a 2 × 2 repeated measures analysis of variance (rmANOVA) with average target distance as the dependent variable. This analysis revealed an effect of block type, *F*(1,15) = 14.08, *p* = .002, η^2^ = .484, but no effect of block number, *F*(1,15) = .985, *p* = .337, and no interaction between block type and block number, *F*(1,15) = .778, *p* = .392. The thick line in Figure 5C shows the group means of the average target distance for all force-on trials and all force-off trials (collapsing across block numbers because there was no main effect of block). The thin gray lines represent individual participants. We found that participants in force-off trials (M = 11.03 cm, SE = 0.36) visited objects that were, on average, .72 cm (SE = .19) further away from the start position than in force-on trials (M = 10.31 cm, SE = .42). These results indicate that although participants tended to search significantly closer to the start position in force-on trials, the effect was small. That is, 0.72 cm is considerably less than the average x or y distance between adjacent targets (3.5 cm).

There are a number of factors that may have contributed to the finding that effort had a significant, albeit small, influence on search behaviour in Exp. 2 but not in Exp. 1. First, participants experienced higher peak forces on the handle of the manipulandum in Exp. 2 compared to Exp. 1. On average, in a given trial, participants in Exp. 2 experienced a peak force of 21.0 N (SE = .4) on the handle whereas the average peak in Exp. 1 was 12.0 N (SE = .5). Second, whereas participants in Exp. 1 only experienced forces during a short duration point to point movement, participants in Exp. 2 experienced these forces over a longer time period when they were visiting and holding the handle over objects during search. Third, whereas movement costs were experienced during the search process itself in Exp. 2, movement costs were experienced after search was completed in Exp. 1 and therefore may have been temporally discounted (Berret & Jean, 2016; Rigoux & Guigon, 2012; Shadmehr, Orban de Xivry, et al., 2010). The greater and longer lasting forces experienced in Exp. 2 may have led to participants becoming fatigued and it has been shown, in several tasks, that fatigue can be a motivating factor in making movement decisions that will reduce effort (Iodice, Calluso, et al., 2017; Iodice, Ferrante, et al., 2017).

### Experiment 3

The previous experiments examined whether movement effort can influence search behaviour during the performance of visual and manual search tasks. Given that movement time costs have been found to influence choice behaviour in humans (Berret & Jean, 2016; Rigoux & Guigon, 2012), we next investigated whether movement time costs can influence search behaviour. In Exp. 3 we used a manual search task that was similar to the task used in Exp. 2 to investigate whether people are sensitive to movement time costs when searching for a target object.

Participants searched a grid containing 4 target objects and 56 distractor objects. To reveal the identity of an object as either a target or distractor, participants had to hold a cursor over its location. In this task participants controlled the speed and direction of cursor motion using a virtual joystick simulated with a robot handle. To manipulate the cost of time, the gain between the excursion of the joystick from its central (resting) position and cursor speed—which dictated the maximum speed of the cursor when the joystick reached its maximum excursion—was varied as a function of the location of the cursor in the search space. More specifically, the maximum speed (or gain) depended on the angle of the cursor relative to the center of the search space. By looking at the location of objects visited, we asked whether search was biased towards regions of space with higher maximum cursor speeds, which we expected would result in lower time costs associated with search.

In slow-left trials, the maximum speed was greatest when the cursor was located to the right of center (0 degrees) and slowest when the cursor was located to the left of center (180 degrees), and these directions were flipped in slow-right trials. The four target objects in a given trial were randomly located such that there was a high probability (p = .9375) that at least one target would be located on a given side (left or right). Thus, participants could, in general, reduce the time required to locate a target object by searching the side of space associated with faster cursor movements. We predicted that search would be biased toward the side of the search space where the cursor speed was highest; i.e., that participants would be sensitive to time costs associated with search.

Each participant experienced two slow-left blocks of trials and two slow-right blocks of trials, with the two block types alternating. The initial block type was counterbalanced across participants. We implemented this block structure given the possibility that any effect of the cursor speed on search behaviour might only emerge after experiencing a number of slow-left and slow-right trials (i.e., that a bias might only emerge after participants learned the relative time costs associated with the two different environments).

## Methods

### Participants

Twelve participants (6 female) between the ages of 18 and 21 years old (M = 18.9) were recruited for this experiment. (See Experiment 1 for sample size considerations.) Participants were required to be right handed, and have normal or corrected-to-normal vision while wearing contacts. All participants were compensated $15 for their participation. Participants provided written informed consent, and after the conclusion of the experiment they were debriefed. The experiment was approved by the Queen’s General Research Ethics Board and complied with the Declaration of Helsinki.

### Apparatus & Stimuli

Participants used the same apparatus as in Exp. 1. The position of the cursor and its velocity were recorded at 1000 Hz. Eye movements were not recorded. The target and distractor objects in this experiment were the same size and appearance as in Exp. 2. As in Exp. 2, these were placed within a circle (radius of 14 cm) and aligned to a grid containing 61 cells. The method for selecting target locations on each trial was the same as that described in Exp. 2, except rather than randomly varying the cursor’s start location on each trial, the start location was always in the cell located in the center of the search grid.

To create a virtual joystick, we simulated, using a very stiff damped spring (6000 Ns/m stiffness, -4 N/m damping), a circular barrier of radius 1 cm around the home position of the handle. Thus, handle movement was limited to 1 cm in any direction. Additionally, a weak damped spring (300 Ns/m stiffness, -1 N/m damping) generated forces on the handle towards the home position. Thus, if no forces were applied to the handle by the participant, this spring brought the handle back to its home location. These two springs allowed the handle of the manipulandum to effectively function as a joystick. The location of the handle while operating as a joystick was ∼20 cm in front of the participant’s chest and in the mid-sagittal plane.

The cursor’s speed depended on the distance and direction of the joystick from its central start position and the current angular location of the cursor according to the following relationship:

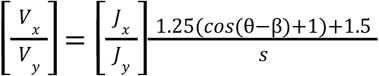

where V_x_ and V_y_ are the x and y cursor velocities in cm/s, J_x_ and J_y_ are the x and y joystick positions in cm, *θ* is the angular position of the cursor, and β is either 0° or 180° in the slow-left and slow-right conditions, respectively. We applied a cosine function to the cursor’s angle in order to allow for a gradual change in cursor speed between the 0° and 180° positions. The perimeter of the blue region in Figure 6A represents the maximum cursor speed (with the joystick at full excursion) as a function of angle in the slow-left condition where the maximum speed is 1.5 cm/s when *θ* = 180° and 4 cm/s when *θ* = 0°.

**Figure 6.**
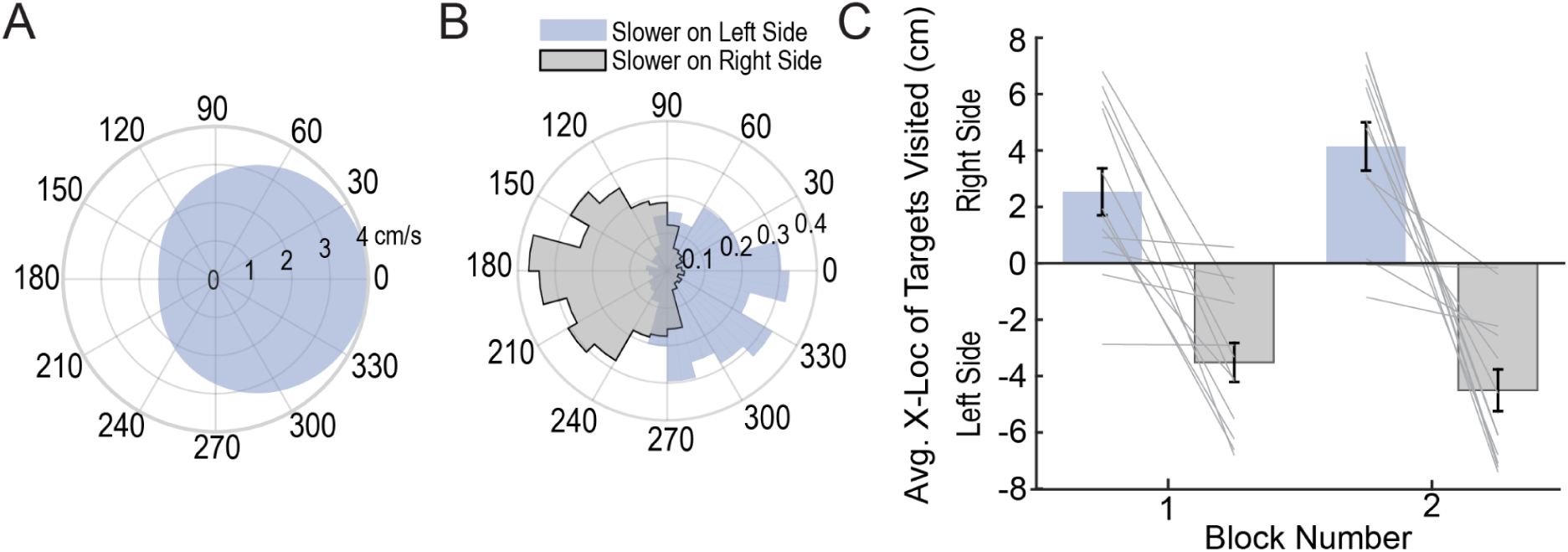
A) Polar plot indicating the relationship between cursor angle and the maximum speed of the cursor (perimeter of the blue shaded area) in the slow-left condition in which the cursor is slowest at 180 degrees (1.5 cm/s) and fastest at 0 degrees (4 cm/s). B) Polar plot showing the probability density, combining all data from all participants, of the cursor’s angular location at visited objects (15° bins). In the slow-left (blue) and slow-right (gray) conditions, participants tended to visit objects on the right and left sides of the search space, respectively. C) Average x-location of objects visited by the cursor in each trial block. Positive values indicate locations to the right of midline. Gray lines represent individual participants. Error bars indicate ±1 standard error.

Participants had to locate one of the four targets after the search objects appeared. In order to reveal the identity of an object the participant had to keep the center of the cursor within 5 mm of the center of the object for 300 ms. The same feedback as in Exp. 2 was given for either successfully locating the target object, or timing out, if the search time exceeded 30s.

### Procedure

Participants completed four blocks of 25 trials starting with either a block of slow-right trials or slow-left trials (counterbalanced across participants), and then alternating between block types. Prior to beginning each block, participants completed a single practice trial, where they had to visit each search object location and reveal its identity with the cursor. (All objects were ‘distractors’ and turned gray when visited.) On this single practice trial, cursor behaviour was the same as in the upcoming block and participants were told that this would be the case. These practice trials were included so that participants had an opportunity to experience and learn the joystick-to-cursor mapping applied to trials within the upcoming block.

## Results and Discussion

We observed no significant difference between slow-left and slow-right trials in terms of search time, *t*(11) = .726, *p* = .483, or the number of objects visited, *t*(11) = .112, *p* = .913. Overall, the average search time was 18.7 s (SE = .48) and the average number of objects visited was 9.9 (SE = .29).

To investigate the influence of movement time on participants’ search behaviour we looked at both the angle and x-location of objects visited across block types. Figure 6B shows a polar probability density plot of the location of objects visited for each block type. It can be seen that participants tended to visit objects on the side associated with faster cursor movements, with most of the visited objects located within 60 degrees of the angle associated with the fastest cursor speed (i.e., a region where the maximum cursor speed was at least 84% of the fastest cursor speed).

To further assess the influence of movement time on search behaviour, for each participant we computed the average x-location of objects visited in each trial and then determined the average for each trial block. Figure 6C shows the average x-location of objects visited in the first and second blocks of slow-left and slow-right trials. To investigate the effects of the block type (slow-left versus slow-right) and block number (first or second), we ran a 2 × 2 repeated measures analysis of variance (rmANOVA) with the average x-location of objects visited as our dependent variable. This analysis revealed an effect of block type, *F*(1,11) = 27.93, *p* < .001, η^2^ = .717, but no effect of block number, *F*(1,11) = .44, *p* = .519, η^2^ = .039. However, there was an interaction between block type and block number, *F*(1,11) = 11.10, *p* = .007, η^2^ = .502. The significant interaction was driven by a larger difference between block types (i.e., slow-left vs. slow-right trials) in the second block of trials (8.65 cm) compared to the first block of trials (6.05 cm). Follow up paired t-tests revealed significant effects of block type in both the first block, *t*(11) = 4.81, *p* = .001, and the second block, *t*(11) = 5.37, *p* < .001, which remain significant when corrected with the Holm-Bonferroni method. Taken together, these results suggest that participants searched in areas associated with higher cursor speed, and hence, lower time costs, and that this bias increased as participants gained experience with the task and became more familiar with the search environment.

The fact that participants’ search was biased to the side of the search space associated with a higher maximum cursor speed, does not necessarily imply that they actually shortened their search times. Specifically, it is possible that participants did not take advantage of the faster cursor speed. To examine the speed at which participants searched, for each participant we calculated the average duration (across all trials) of cursor movements between search objects located on the fast side of the search space and between search objects located on the slow side (we did not consider movements between objects located on opposite sides of midline). We found that movement durations were significantly shorter, *t*(11) = 10.09, *p* < .001, for cursor movements between fast side objects (M = 1.09 s, SE = .02) than between slow side objects (M = 1.69 s, SE = .06). This result indicates that participants exploited the variable cursor speed to reduce their search times.

The results of this experiment are consistent with previous work that has found that people incorporate kinematic factors such as movement time and effort, as well as as object size and distance (which influence movement time), when deciding between movement options (Cos et al., 2012, 2014; Michalski et al., 2020). Our current results suggest that the influence of movement time costs extends to decisions involved in search, which are traditionally considered to be more cognitive in nature.

### Experiment 4

The results of Exp. 3 suggest that movement time costs can influence behaviour in a manual search task. In Exp. 4, we investigated whether movement time costs can also influence visual search behaviour. The findings from Exp. 3 showed that the bias of cursor speed on search location increased with experience. This change in bias suggests that experience moving in the search environment was required for participants to fully learn about the time costs and apply this knowledge during search. Therefore, before testing participants on a visual search task that incorporated movement time costs, we trained them on a manual search version of the task so that they had the opportunity to learn about the costs associated with moving in different regions of the search space.

In Exp. 4, participants performed the manual search task described in Exp. 3 as well as a visual search version of the task in which they had to move the cursor to a target after a target was found (which required foveating the target object). Participants first completed a block of manual search trials with the slow side (left or right) counterbalanced across participants, and then completed a block of visual search trials that had the same mapping between maximum cursor speed and side of space. Participants then completed a block of manual search trials with the slow side on the other side of space, followed by a block of visual search trials where, again, the mapping between speed and side of space was preserved from the previous manual search trial block. Note that participants were explicitly told prior to each block of visual search trials that the mapping would be the same as in the block of manual search trials they had just experienced.

We predicted that participants would learn the relationship between cursor speed and angle during the manual search trials and demonstrate a manual search bias towards the side of space with faster cursor speeds. We further predicted that they would subsequently demonstrate a bias towards searching the side of space associated with faster cursor speeds in the following visual search trials.

## Methods

### Participants

Twelve participants (8 female) between the ages of 18 and 22 years old (M = 19.8) were recruited for this experiment. (See Experiment 1 for sample size considerations.) Participants were required to be right handed, and have normal or corrected-to-normal vision while wearing contacts. All participants were compensated $15 for their participation. Participants provided written informed consent, and after the conclusion of the experiment they were debriefed. The experiment was approved by the Queen’s General Research Ethics Board and complied with the Declaration of Helsinki.

### Apparatus & Stimuli

Participants used the same apparatus used in Exp. 1 (see Fig. 1B). Search objects in the manual search trials were gray squares (width 1 cm; ∼1.6 degrees visual angle) and the participant could identify a square as either a target or a distractor by holding the cursor over its location. Objects in the visual search trials were split colour squares (width 1 cm) with one half pink and the other half blue (see Figure 7A). The target objects had the opposite colour arrangement to the distractor objects. The position of the cursor was recorded at 1000 Hz. Gaze data was collected at a rate of 500 Hz using an infrared eye tracker (Eyelink 1000, SR Research, Ottawa, ON, Canada). A chin rest was used to limit head motion during the experiment.

**Figure 7.**
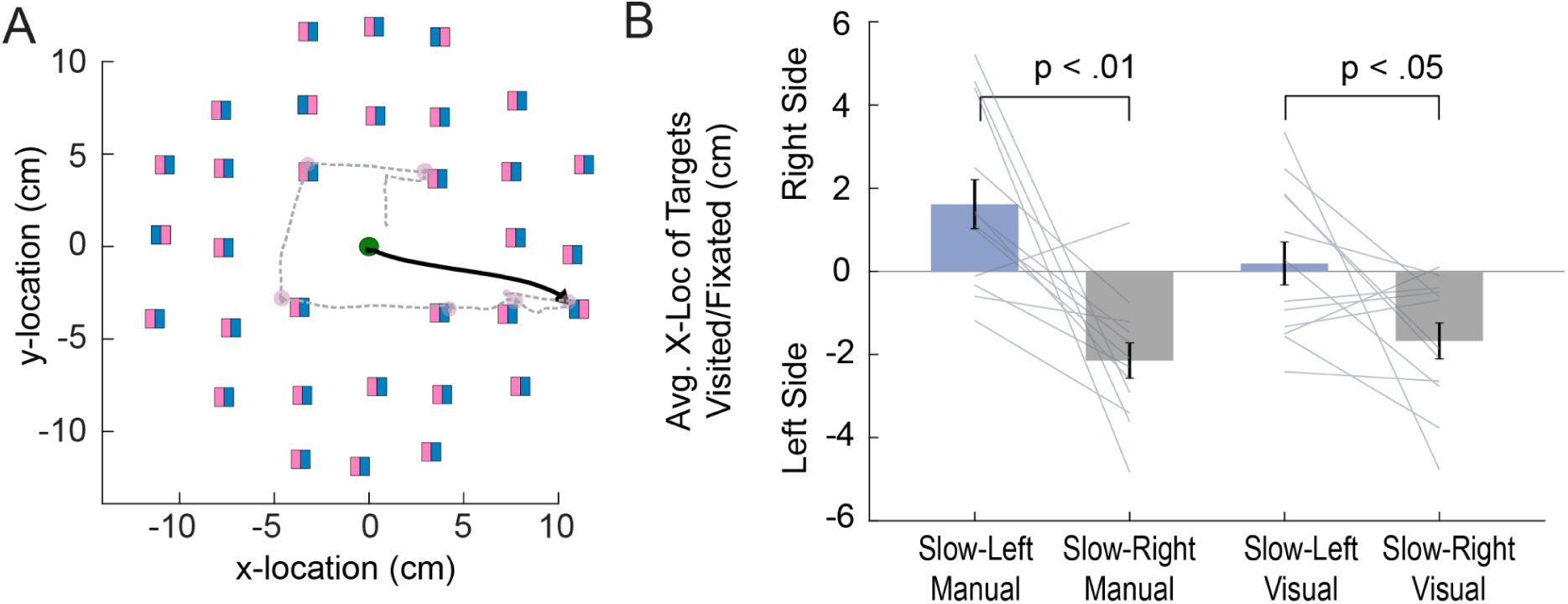
A) In each trial, 32 objects were presented on the screen located in cells, each 3.8 × 3.8 cm in size. The object position within each cell was randomly jittered. The 32 objects included four target objects (pink on the right side), and 28 distractor objects (pink on the left side). The gray dashed line shows an example gaze path for a slow-left visual search trial starting from the center and ending at a target object on the right side of the screen. The solid black trace shows the cursor path taken to the target object. B) Average x-location of objects visited by the cursor in manual search trials (left bars) and fixated in visual search trials (right bars). Positive values indicate locations to the right of midline. Gray lines represent individual participants. Error bars indicate ±1 standard error.

Target and distractor objects were presented in a circular search area with a radius of 11.4 cm (see Figure 7A). Within this circle, we arranged 32 search objects by aligning them to a grid containing 33 cells, with the extra cell containing the start position at the center. The size of each cell of the grid was 3.8 × 3.8 cm. There were four search targets in each trial and the method for selecting target locations was the same as that described in Exp. 3.

Manual search trials proceeded in the same manner as described in Exp. 3. For visual search trials, participants were instructed to move the cursor from the start position to the target, once they visually located a target, and hold the cursor at that location to end the trial. The hold criteria for the target in visual search trials was identical to the hold criteria for revealing a search object’s identity in the manual trials, which was fully described in the methods for Exp. 3. When moving, the search cursor behaved in an identical manner to manual search trials in the previous trial block. That is, the fast and slow sides of the search space (with maximum cursor speeds of 4 and 1.5 cm/s respectively) were the same as in the preceding block of manual search trials. The feedback given for successfully locating the target object, or timing out, was the same as in Exps. 2 and 3. However we modified the maximum search time (i.e., time out duration) for both manual and visual search trials, extending it from 30 to 60 s since a few manual search trials in Exp. 3 timed out. In manual search trials, the cursor had to be located at the center object at the start of the trial and in visual search trials, participants had to fixate the center object at the start of the trials.

### Procedure

After an initial eye tracking calibration procedure was performed, participants completed four blocks of 25 trials. They first performed a block of manual search trials and then a block of visual search trials with the slow side either on the left or right (counterbalanced across participants) in both blocks. Following these two blocks, they performed a block of manual search trials and then a block of visual search trials with the slow side on the opposite side. After completing a block of manual search trials, participants were explicitly informed that the relationship between cursor location and speed would be the same on the subsequent block of visual search trials. Prior to beginning each manual search block, participants completed a practice trial, as in Exp. 3, where the cursor’s behaviour was the same as in the upcoming block of trials. We modified this trial slightly from the previous experiment, reducing the number of objects participants had to visit to 16. These practice trials were included so that participants had an opportunity to experience and learn the joystick-to-cursor mapping applied to trials in the upcoming block of manual trials.

### Data Analysis

For manual trials, we extracted the sequence of objects visited by the cursor during the search period, and in visual search trials, after eliminating blinks and low-pass filtering the raw gaze signal, we extracted the fixation locations for each trial during the search period. In each trial, we assigned each fixation (except the first) to the closest search object and used the location of those objects as a measure of where participants searched. This allowed us to make a direct comparison between where participants searched in visual and manual search trials.

## Results & Discussion

For manual search blocks, there was no significant difference in search time between slow-right (M = 15.6 s, SE = .9) and slow-left (M = 15.8 s, SE = .7) trials, *t*(11) = .161, *p* = .875. Likewise, for visual search blocks, there was no significant difference in search time between slow-right (M = 5.8 s, SE = .2) and slow-left (M = 6.1 s, SE = .2) trials, *t*(11) = 1.34, *p* = .207.

We examined, for each participant, the average x location of objects contacted by the cursor in manual search trials or fixated in visual search trials. Figure 7B shows, for both manual and visual search trials, the average x-location of visited objects for slow left and slow right trials. In manual search trials, on average search was biased toward the ‘fast’ side of the search space. In visual search trials, search was biased toward the fast side when that side was on the left (slow-right trials) but no clear bias was observed when the fast side was on the right (slow-left trials). One explanation for this pattern of results is that, in visual search, there is both a bias towards search on the left and a bias towards searching on the fast side.

To examine the influence of block type on manual and visual search behaviour, we conducted a 2 (block type: slow-left, slow-right) × 2 (search type: manual, visual) rmANOVA. The analysis revealed a significant effect of block type, *F*(1,11) = 14.70, *p* = .003, η^2^ = .572, but not search type, *F*(1,11) = 4.06, *p* = .069, η^2^ = .270, while the interaction between search type and block type approached significance, *F*(1,11) = 4.61, *p* = .055, η^2^ = .295. Follow up paired t-tests revealed significant differences between block type for both manual, *t*(11) = 4.08, *p* = .002, and visual, *t*(11) = 2.37, *p* = .037, search trials, which remained significant when corrected with the Holm-Bonferroni method (see Figure 7B).

Our finding that manual search is biased by movement time costs (associated with cursor speed) replicates the results obtained in Exp. 3. Our finding that visual search can also be biased by cursor speed can be contrasted with the results of Exp. 1, in which movement effort cost did not influence visual search. This suggests that time cost may be more aversive than effort cost, at least as implemented in our experiments.

## General Discussion

Converging evidence from a number of studies suggests that movement costs, such as effort or time, can influence the decisions we make during the performance of action tasks (Bakker et al., 2017; Burk et al., 2014; Cos et al., 2011, 2012, 2014; Michalski et al., 2020; Morel et al., 2017). In this paper we examined how movement costs incurred either in the act of searching (manual search), or when reaching to a target once it is visually located (visual search), influence search behaviour. Across four experiments we tested whether search behaviour is biased by motor costs by varying effort or time costs across the search environment. We were interested in whether participants would take these costs into account while searching such that they would bias their search to areas of the search environment that reduced movement effort or movement duration.

In Exp. 1, we examined whether search behaviour in visual search is biased by motor effort, and failed to find an effect. In Exp. 2, we asked whether search behaviour in a manual search task can be biased by motor effort and did demonstrate a small effect of effort. In Exp. 3, we tested the influence of movement time costs on search behaviour in a manual search task, and found a strong effect. Finally, in Exp. 4, we examined whether movement time costs bias search behaviour in a visual search task, and found that these costs did influence search behaviour. To summarize, we demonstrated that effort-based costs have an overall weak influence on human search behaviour, with a small influence on manual search behaviour but none on visual search behaviour. In contrast, time-based costs appear to have a strong influence on manual search but also influence visual search. Thus, the current study shows that movement time and effort costs can influence human search behaviour, at least in some contexts.

Previous work has shown that movement costs are factored into human decision making across a variety of tasks. For example, it has been shown that movement costs can influence the choice of hand (left or right) used to perform a target reaching task (Bakker et al., 2017), the choice of which target to reach towards (Cos et al., 2014; Morel et al., 2017), the extent to which people opt to rely on memory during search (Kit et al., 2014; Li et al., 2016, 2018), and perceptual judgement tasks (Burk et al., 2014). The novel contribution of the current work is the demonstration that movement costs can influence both visual search when participants reach to the target object after locating it, and manual search when participants make reaching movements to objects to determine which is the target.

As noted in the introduction to this paper, it has been suggested that, under conditions of target uncertainty, the brain prepares multiple actions in parallel, before selecting one of them to execute (Cisek, 2007; Cisek & Kalaska, 2005; Gallivan et al., 2015, 2017, 2018; Gallivan, Logan, et al., 2016; Klaes et al., 2011; Pastor-Bernier & Cisek, 2011; Song & Nakayama, 2009; Stewart et al., 2014; Thura & Cisek, 2014; Wispinski et al., 2020). Planning competing potential actions, in parallel, may provide a mechanism through which the motor costs associated with these actions can be taken into account when deciding which one to execute (Cisek, 2006). Note that this parallel planning hypothesis is related to the idea of affordances whereby objects in the environment automatically evoke action representations (Gibson, 1979). For example, it has been shown that visual displays of cups can automatically prime reach and grasp movements (Handy et al., 2003; Masson et al., 2011).

Most of the neurophysiological and behaviour work supporting the ‘parallel planning’ hypothesis has used tasks in which there are two competing targets (e.g., Cisek & Kalaska, 2005; Gallivan et al., 2017). Although a few of these studies have argued that we can plan more than two competing potential movements (Chapman et al., 2010; Stewart et al., 2013), it seems highly improbable that in our search tasks, involving up to 60 objects, people simultaneously plan reaching movements for all objects before deciding which one to move their hand (manual search) or gaze (visual search) to next. However, given that the search behaviour we observed typically involved sequential eye or hand movements to nearby objects, it is conceivable that the brain prepares parallel reach plans for a small set of objects in close vicinity of the current object being inspected. It is also possible that in manual search, the brain may plan, in parallel, a short sequence of forthcoming actions. Several studies have provided evidence that when planning a short sequence of manual actions, the goals of each action are represented, in parallel, in sensorimotor areas (Andersen & Cui, 2009; Baldauf et al., 2008; Gallivan, Johnsrude, et al., 2016). Moreover, a recent study examining a task in which participants made reaching movements to ‘harvest’ as many target objects as possible within a short time period, decisions about which target to move to next took into account the cost of moving not only to that target but also future targets (Diamond et al., 2017).

Although we propose that search behaviour in our tasks was influenced by biomechanical effort and movement time, there may be other factors that could have influenced participant behaviour. One type of movement cost that may have played a role in influencing search behaviour is metabolic fatigue. In Exp. 1 we failed to note any significant influence of the resistive force on search behaviour, despite previous work that found that such forces have the capacity to bias perceptual decision making (Hagura et al., 2017). One possible source of this discrepancy could be the amount of fatigue participants experienced through exposure to forces. In the test phase of Exp. 1, resistive forces were only experienced during the reach at the end of the trial and participants performed a total of 180 trials. In contrast, participants in the Hagura et al. (2017) study completed nearly five times as many trials with loads applied. It is possible that in their experiment, participants became fatigued, which could have made forces more salient. The significant influence of effort on search behaviour we noted in Exp. 2 could have also resulted from fatigue, owing to the larger and greater duration of the applied forces, when compared to Exp. 1. As participants become more metabolically fatigued from reaching, there might have been a corresponding increase in the penalty of effort costs, leading to a greater influence of expected effort on movement decisions (Iodice, Calluso, et al., 2017).

It is possible that search behaviour in everyday search tasks in familiar environments is more strongly influenced by movement costs than in our laboratory tasks. Consider, for example, searching for your cell phone at home. In this case, you have ample opportunity to learn the costs associated with moving in the search environment. In contrast, participants in our experiments had no previous experience with the cost structure of the search environment. Nevertheless, it is clear from Exps. 2, 3 and 4 that participants were able to learn the cost structure of the search when performing manual search and, moreover, that this learning could transfer to visual search.

In our tasks, the forces we applied to the handle of the manipulandum, and the speed limits we placed on cursor motion, were quite artificial manipulations of movement effort and movement time, respectively. It is possible that in more ecological tasks, where movement costs are manipulated in a less artificial manner, that we would see a larger influence of movement costs on participant behaviour. In future work, we plan to examine other ways of manipulating movement effort and time costs that may be more naturalistic. For example, we could place an obstacle in the search space during a visual search task in which participants reach to a target once it is found, and ask if search is biased away from the regions where reaching to a target would require moving around the obstacle. Given the presence of obstacles in our everyday environment it is possible that participants can readily take such movement costs into account when making decisions about where to search.

The current findings show the importance of factoring in movement costs into our understanding of real-world search behaviour. Future studies would benefit from more closely examining the relationship between rewards, costs, and memory limitations that may influence real world search and other action tasks. Our study adds to the growing body of evidence that factors impacting on the motor system, which is often viewed as the final output step in producing behaviour, can also influence perceptual and cognitive processes related to decision-making during task performance.

